# Impaired Visuospatial Working Memory but Preserved Attentional Control in Bipolar Disorder

**DOI:** 10.1101/2024.01.04.574011

**Authors:** Catherine V. Barnes-Scheufler, Lara Rösler, Carmen Schiweck, Benjamin Peters, Silke Matura, Jutta S. Mayer, Sarah Kittel-Schneider, Michael Schaum, Andreas Reif, Michael Wibral, Robert A. Bittner

**Author notes:** Corresponding authors. Goethe University Frankfurt, University Hospital, Department of Psychiatry, Psychosomatic Medicine and Psychotherapy, Heinrich-Hoffmann-Str. 10, D-60528 Frankfurt am Main, Germany.

## Abstract

**BACKGROUND:** Persistent deficits in working memory (WM) and attention have considerable clinical and functional impact in people with bipolar disorder (PBD). Understanding the neurocognitive underpinnings of these interacting cognitive constructs might facilitate the discovery of more effective pro-cognitive interventions. Therefore, we employed a paradigm designed for jointly studying attentional control and WM encoding.

**METHODS:** We used a visuospatial change-detection task using four Gabor Patches with differing orientations in 63 euthymic PBD and 76 healthy controls (HCS), which investigated attentional competition during WM encoding. To manipulate bottom-up attention using stimulus salience, two Gabor patches flickered, which were designated as either targets or distractors. To manipulate top-down attention, the Gabor patches were preceded by either a predictive or a non-predictive cue for the target locations.

**RESULTS:** Across all task conditions, PBD stored significantly less information in visual WM than HCS (significant effect of group). However, we observed no significant group by salience or group by cue interactions. This indicates that impaired WM was not caused by deficits in attentional control.

**CONCLUSIONS:** Our results imply that while WM is disturbed in PBD, attentional prioritization of salient targets and distractors as well as the utilization of external top-down cues were not compromised. Consequently, the control of attentional selection appears be intact. These findings provide important constraints for models of WM dysfunction in PBD by indicating that later stages of WM encoding are likely primarily impaired. We also demonstrate that selective attention is not among the main sources of cognitive impairment in PBD.

## INTRODUCTION

Cognitive impairment in persons with bipolar disorder (PBD) is a core feature persisting during euthymia phases (Quraishi & Frangou, 2002). It is a key predictor for quality of life (Green, 2006), and for important measures of clinical and functional outcome including recurrence of affective episodes and occupational disability (Sachs, Berg, Jagsch, Lenz, & Erfurth, 2020). However, sufficiently effective pro-cognitive interventions for PBD remain elusive (Miskowiak et al., 2022), mainly because we lack a thorough understanding of the underlying neurocognitive architecture (Tamura et al., 2022).

Impaired information processing in bipolar disorder (BD) has been linked to its neurodevelopmental origins (Bortolato, Miskowiak, Köhler, Vieta, & Carvalho, 2015). Within the proposed neurodevelopmental continuum, which also encompasses intellectual disability, autism spectrum disorders, schizophrenia, and attention-deficit/hyperactivity disorder (ADHD) (Owen & O’Donovan, 2017), BD is considered to have the lowest degree of neurodevelopmental and cognitive disturbances, with a unique cognitive profile (Valli, Fabbri, & Young, 2019).

Both working memory (WM) and attention are regarded as central cognitive domains for transdiagnostic studies of impaired information processing (Insel et al., 2010), and deficits in both domains remain stable over time in PBD (Kjærstad, Søhol, Vinberg, Kessing, & Miskowiak, 2023). Specifically, WM dysfunction is a crucial deficit in both PBD and schizophrenia with stronger impairment observed in schizophrenia (Barnes-Scheufler et al., 2021). It remains the most consistently reported cognitive deficit in PBD, making it a key target for pro-cognitive interventions in PBD (Miskowiak et al., 2018). Similar levels of WM impairment have been observed in both type I and type II BD (Bora, 2018; Mayer & Park, 2012), but PBD with a history of psychosis appear to be more affected than other PBD (Bora, 2018). WM capacity is reduced even in euthymic PBD (Barnes-Scheufler et al., 2021; Mayer & Park, 2012). Importantly, functional neuroimaging studies indicate disturbances during WM encoding in areas closely involved in both WM and attention (Huang et al., 2019; Mayer et al., 2007), and a further exacerbation of aberrant activity following affective phases (Macoveanu et al., 2021). Deficits in visual attention are present in unaffected first-degree relatives (Bora, Yucel, & Pantelis, 2009), however impairment in sustained attention does not seem to be related to aberrant WM performance (Harmer, Clark, Grayson, & Goodwin, 2002).

Although some aspects of attention have been investigated in PBD, studies investigating bottom-up and top-down attention remain scarce. Yet, these processes are of particular importance given their close connections to WM (Oberauer, 2019). Top-down attention influences visual WM (VWM) capacity because of its close involvement in selecting relevant information during VWM encoding (Vogel, McCollough, & Machizawa, 2005). This constitutes an important candidate mechanism for reduced WM capacity in BD, especially given the evidence for impaired WM encoding (Barnes-Scheufler et al., 2021; Huang et al., 2019). However, the contribution of attentional dysfunction to impaired WM encoding in BD remains unclear.

The successful encoding of relevant information into VWM and the suppression of irrelevant information depends on interactions between bottom-up and top-down attention, which are driven by stimulus salience and stimulus relevance respectively. The resolution of attentional competition elicited by both stimulus features determines the probability of encoding an object into VWM (Constant & Liesefeld, 2021; Liesefeld, Liesefeld, Sauseng, Jacob, & Müller, 2020). Importantly, two top-down control processes support the selection of relevant information when multiple items compete for attention: the control of selection which assists in the identification of relevant information, and the implementation of selection which differentially processes relevant and irrelevant information (Desimone & Duncan, 1995; Luck & Gold, 2008). Neural computations during attentional competition assign a distinct priority to each stimulus, based on both its salience and its behavioral relevance (Fecteau & Munoz, 2006). An attentional set aids the control of attentional selection by guiding top-attention to the most relevant information based on current goals (Gaspelin & Luck, 2018a). According to the signal suppression hypothesis, this mechanism also controls the active suppression of automatic attentional capture by visually salient distractors via inhibitory mechanisms (Gaspelin & Luck, 2018b). The parallelized up- and down-weighting of stimuli is essential for the generation of a priority map and the implementation of attentional selection (Gaspelin & Luck, 2018b).

We used these findings from cognitive neuroscience as a framework for our investigation of top-down and bottom-up attention during VWM encoding in PBD. To this end, we employed a visuospatial change detection task with an encoding array containing an equal number of salient (flickering) and non-salient (non-flickering) Gabor patches with different orientations. Depending on the specific task condition, either the salient or non-salient Gabor patches would be most relevant and probed preferentially (Constant & Kerzel, 2025; Constant & Liesefeld, 2023; Emrich, Lockhart, & Al-Aidroos, 2017; Lockhart, Dube, MacDonald, Al-Aidroos, & Emrich, 2024; Ravizza & Conn, 2022). The encoding array was preceded by either a predictive external cue indicating the location of goal-relevant stimuli, or a non-predictive external cue providing no such information.

Previously, we utilized the same paradigm to study people with schizophrenia (PSZ) (Barnes-Scheufler et al., 2023). We observed a significant VWM deficit in PSZ, with less information encoded by PSZ across all conditions. In addition, we reported significant main effects of cue and salience, as well as interactions of group by cue and group by salience indicating group differences in top-down and bottom-up processing. Post-hoc results showed that PSZ stored significantly more non-flickering information with the help of a top-down predictive cue, mirroring an earlier report of successful utilization of top-down cues in PSZ (Gold et al., 2006). Furthermore, in the non-predictive cue conditions, PSZ stored significantly more flickering than non-flickering information-indicative of a bottom-up attentional bias in PSZ (Hahn et al., 2010).

In line with the neurodevelopmental continuum model and the clear evidence for overlapping cognitive deficits in schizophrenia and BD (Bora & Pantelis, 2015), we expected to observe a pattern of similar, yet less pronounced disturbances in PBD. Specifically, we predicted an overall deficit in the amount of information stored in VWM in PBD as a main group effect, and interactions of group by salience and group by cue to indicate deficits in top-down and bottom-up attentional processing respectively.

## METHODS AND MATERIALS

### PARTICIPANTS

63 PBD and 76 healthy control subjects (HCS) participated in this study. Patients were recruited from outpatient facilities in and surrounding Frankfurt am Main, Germany. HCS were recruited via online and printed advertisements. 53 HCS were already part of the control group for our previous study investigating PSZ (Barnes-Scheufler et al., 2023). Diagnoses of all patients were established according to DSM-5 criteria involving a clinical interview and careful chart review of a trained psychiatrist. The Young Mania Rating Scale (YMRS) (Young, Biggs, Ziegler, & Meyer, 1978) and Montgomery-Åsberg Depression Rating Scale (MADRS) (Montgomery & Åsberg, 1979) were utilized to establish euthymic mood state in PBD - those with YMRS values of ≥ 11 or MADRS total scores of ≥ 11 were excluded from our analysis. All PBD were on stable medication for at least one month at the time of the study.

No history of any psychiatric disorder or family history of psychiatric disorder in first-degree relatives were reported in HCS. No prior illicit drug use within the past six months and no lifetime history of neurological illness were reported in any participants. All participants reported normal or corrected-to-normal vision, and no color blindness.

Groups did not differ in age, sex, years of education, parental years of education or premorbid IQ, as assessed by the German Mehrfachwahl-Wortschatz-Intelligenz Test (Table 1) (Lehrl, Merz, Burkhard, & Fischer, 2005). Parental years of education was quantified by using the highest value of either parent. The ethics committee of the University Hospital Frankfurt approved all study procedures. Participants provided written informed consent after receiving a complete description of the study and adequate time for questions.

**Table 1.** Values are mean or n. All statistics reported are two-tailed. Abbreviations: HCS = healthy control subjects, PBD = people with bipolar disorder, PBD I = type 1, PBD II = type 2, HPS+= with a history of psychosis, HPS-= without a history of psychosis, YMRS = Young Mania Rating Scale, MADRS = Montgomery-Åsberg Depression Rating Scale, SGA = second-generation antipsychotics, df = degrees of freedom.

### WORKING MEMORY TASK

A visuospatial change detection task utilizing Gabor patches (Figure 1) was implemented on a personal computer using Presentation software version 14.9 (www.neurobs.com). Stimuli were presented on a grey background (RGB values: 191, 191, 191) in a dimly lit room with a viewing distance of approximately 60 cm.

**Figure 1.**
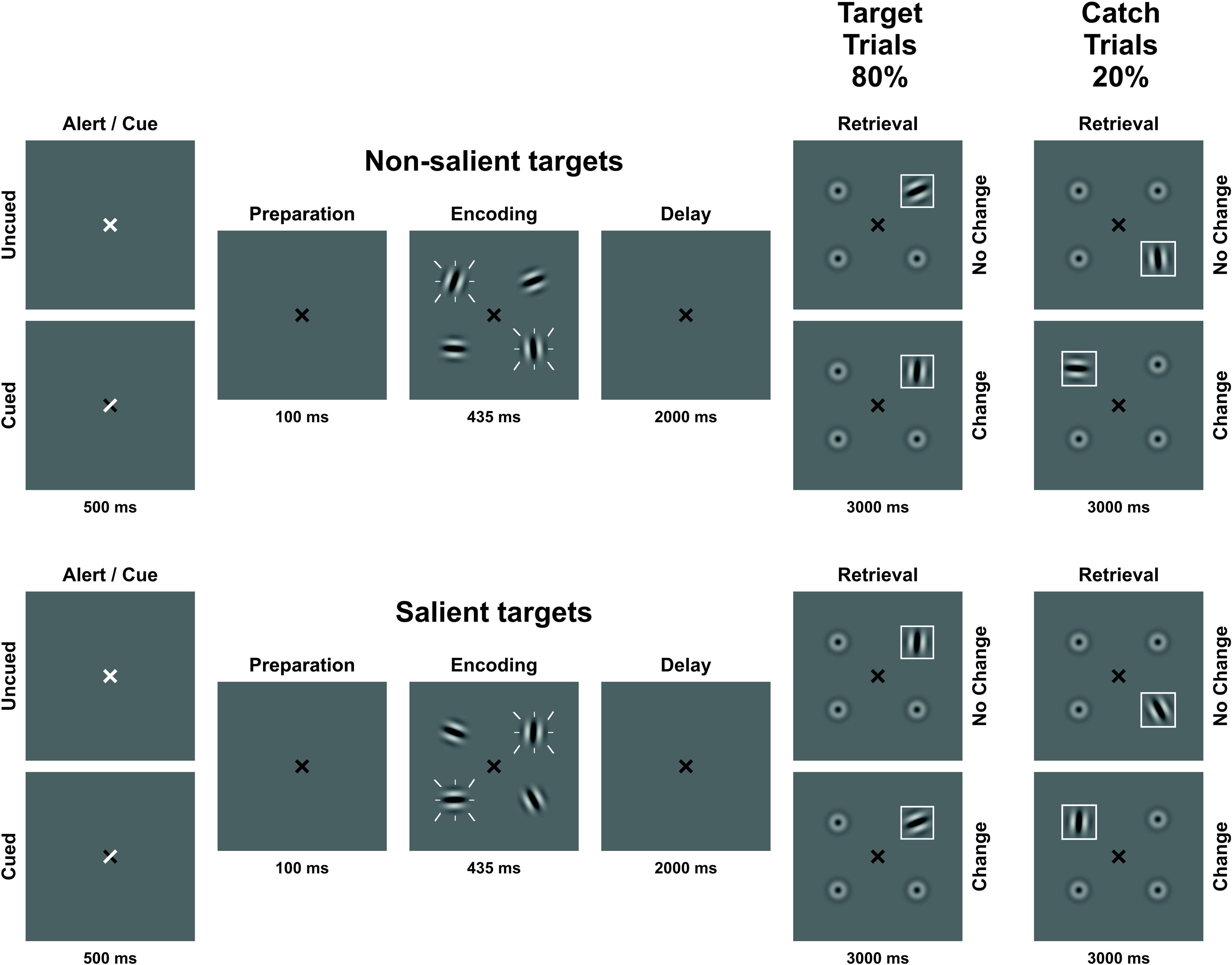
Visual change detection task with four conditions; flickering/predictive cue, flickering/non-predictive cue, non-flickering/predictive cue, non-flickering/non-predictive cue. Flickering is indicated by white dashes around stimuli. The set size of four items was kept constant. In 80% of trials, a designated target stimulus was probed during retrieval (target trials). In 20% of trials, a distractor was probed during retrieval (catch trials).

Gabor patches had a visual angle of 0.96° each and were placed at four fixed locations equally spaced on an imaginary circle (visual angle of the diameter: 3.82°). A black fixation cross (visual angle: 0.48°) was presented at the center of the screen throughout the experiment. At the start of each trial, either a predictive or a non-predictive external cue was presented by either partially (predictive cue) or entirely (non-predictive cue) turning white the fixation cross for 300 ms. For predictive cues, the white arms of the fixation cross indicated the future locations of the two most relevant Gabor patches. For non-predictive cues, the entire fixation cross turned white. After a 300 ms preparation interval, the encoding array consisting of four Gabor patches with different orientations was displayed for 400 ms. To manipulate visual salience, two of the four patches flickered at a frequency of 7.5 Hz. To manipulate stimulus relevance, participants received instructions whether the flickering or non-flickering stimuli would be most goal-relevant and thus probed preferentially, and whether a predictive or non-predictive cue would be displayed, resulting in four conditions: flickering-bias/predictive cue, flickering-bias/non-predictive cue, non-flickering-bias/predictive cue, non-flickering-bias/non-predictive cue.

The delay phase lasted for 2000 ms on average, with a jitter of +/- 250 ms. The retrieval array was displayed for 3000 ms consisting of one Gabor patch surrounded by a white frame at the probed location and three blurred out Gabor patches. Probed locations were randomized but counterbalanced. Participants had to indicate by button press within 3000 ms, if the orientation of the framed Gabor patch was identical to or different from the Gabor patch displayed at the same location in the encoding array. A minimum Gabor patch rotation of 45° indicated a change in orientation.

Both goal-relevant target and goal-irrelevant distractor stimuli were probed during retrieval with 80% and 20% probability, respectively. Probing goal-irrelevant stimuli using catch trials (Figure S1) allowed us to assess the efficiency of attentional prioritization operationalized as the difference of information stored in VWM between target and catch trials. Importantly, predictive cues always indicated the future locations of the target Gabor patches, even in catch trials. A total of 400 trials were presented, 100 for each condition, divided into eight blocks, counterbalanced in order across participants and groups. Each block began with four practice trials to ensure familiarization with the current instructions.

We also employed a 60-trial canonical visual change detection task (Barnes-Scheufler et al., 2021) to obtain an independent estimate of VWM capacity in order to investigate a possible relationship between attentional prioritization and WM capacity (see supplementary materials).

### ANALYSIS OF BEHAVIORAL DATA

We applied the same main statistical analyses used for studying patients with schizophrenia (Barnes-Scheufler et al., 2023). We quantified the amount of information stored in VWM using Cowan’s K, where K = (hit rate + correct rejection rate − 1) × memory set size (Cowan, 2001). Because participants were instructed to encode two Gabor patches in VWM, a set size of two was assumed (Gold et al., 2006). Statistical analyses were conducted using SPSS (IBM) Version 22, and R Version 4.3.1 (www.r-project.org). We created a cutoff value of performance by using a binomial expansion on the 400 trials. It was determined that an accuracy of 56% had a cumulative probability of P(X > x) 0.009, i.e., the probability of getting 224 correct responses or more by chance was less than 1%. Accordingly, we excluded the participants with an accuracy below 56% (n = 5, all PBD). We observed a normal distribution of our data based on visual inspection of QQ-plots and Levene’s Test for all four conditions (*p* = 0.157, 0.831, 0.468, 0.688).

We based our main analysis solely on target trials, to study the differential processing of goal-relevant information. The main analysis consisted of a linear mixed-model (LMM) estimated using ML and nloptwrap optimizer with R to predict Cowan’s K with group, salience, cue and age (Formula: score ∼ Salience * Cue * Group + Age) and subject as a random effect (formula: ∼1 | Subject). This task design was chosen due to the mixed conditions of our study design and therefore repeated measures of the dependent variable. We used an ANOVA function (α = 0.05) to obtain estimates of the LMM. Post-hoc contrasts of the LMM were computed using the Kenward-Roger degrees-of-freedom method with independent sample t-tests adjusted with the Tukey method. The t-tests (α = 0.05) were controlled for age, where age was fixed at 40.5 years as determined by the function. Additionally, we relied on Bayes Factors BF1/BF0 calculated using the R library (BayesFactor) for interpreting our results.

To investigate possible influences of manic and/or depressive symptoms, we correlated overall Cowan’s K of target trials across all four conditions with YMRS and MADRS scores in PBD using two-tailed Spearman correlations. To investigate possible differences in attentional prioritization, independent 2-tailed t-tests were conducted between groups for each condition (Cowan’s K for target trials minus Cowan’s K for catch trials). We assessed handedness as a continuous variable using the Edinburgh Handedness Inventory (Oldfield, 1971), and compared group differences using an independent 2-tailed t-test. Spearman Correlations (two-tailed) were used to investigate possible relationships between gender and overall target Cowan’s K in both groups. Gender was coded as being 0 for female and 1 for male, allowing a ranking system for comparisons.

We correlated attentional prioritization with WM capacity (Pashler’s K), and target Cowan’s K measures with WM capacity using two-tailed Spearman correlations in each condition in each group to investigate WM capacity as a possible limiting factor for attentional prioritization. 95% confidence intervals (CI) were utilized as reliability estimates for the correlational tests, and estimation of standard error is based on the formula proposed by Fieller, Hartley, and Pearson (Fieller, Hartley, & Pearson, 1957).

## RESULTS

### AMOUNT OF INFORMATION STORED IN VWM

We observed a significant main effect of group (F(139) = 8.14, *p* = 0.005; Table 2). For target trials, HCS (65% female) stored 1.25 items across all conditions, corresponding to a performance accuracy of 81% (Figure 2). PBD (75% female) stored on average 1.05 items, with an accuracy of 76%. HCS stored significantly more information into VWM in all four target conditions: flickering-bias/predictive cue (mean = 1.29), flickering-bias/non-predictive cue (mean = 1.27), non-flickering-bias/predictive cue (mean = 1.26), non-flickering-bias/non-predictive cue (mean = 1.18) than PBD: flickering-bias/predictive cue (mean = 1.10, t(253) = 2.35, *p* = 0.019), flickering-bias/non-predictive cue (mean = 1.09, t(253) = 2.23, *p* = 0.027), non-flickering-bias/predictive cue (mean = 1.09, t(253) = 1.99, *p* = 0.048), non-flickering-bias/non-predictive cue (mean = 0.94, t(253) = 3.10, *p* = 0.002). There were no significant correlations between YMRS and overall Cowan’s K for target trials (r_s_ = 0.096, *p* = 0.453, 95% CI [-0.163, 0.343]), as well as MADRAS and overall target Cowan’s K (r_s_ = -0.014, *p* = 0.912, 95% CI [-0.268, 0.242]). Furthermore, there were no effects of medication on Cowan’s K (Supplementary Materials). The covariate of age had a significant effect (F(139) = 21.55, *p* < 0.001). Additionally, we observed a significant interaction of salience and cue (F(417) =1.10, *p* = 0.018). The model’s total explanatory power was substantial (conditional R^2^ = 0.70), and the part related to the fixed effects alone (marginal R^2^) was 0.17, indicating a better fit when random effects were included (Barton & Barton, 2015). We did not observe a group difference in handedness (t_137_ = -0.69, *p* = 0.490). We did not find a correlation between gender and overall Cowan’s K for target trials in PBD (*r_s_* = 0.058, *p* = 0.651, 95% CI [-0.200, 0.308]), or in HCS (*r_s_* = -0.006, *p* = 0.961, 95% CI [-0.237, 0.227]).

**Figure 2.**
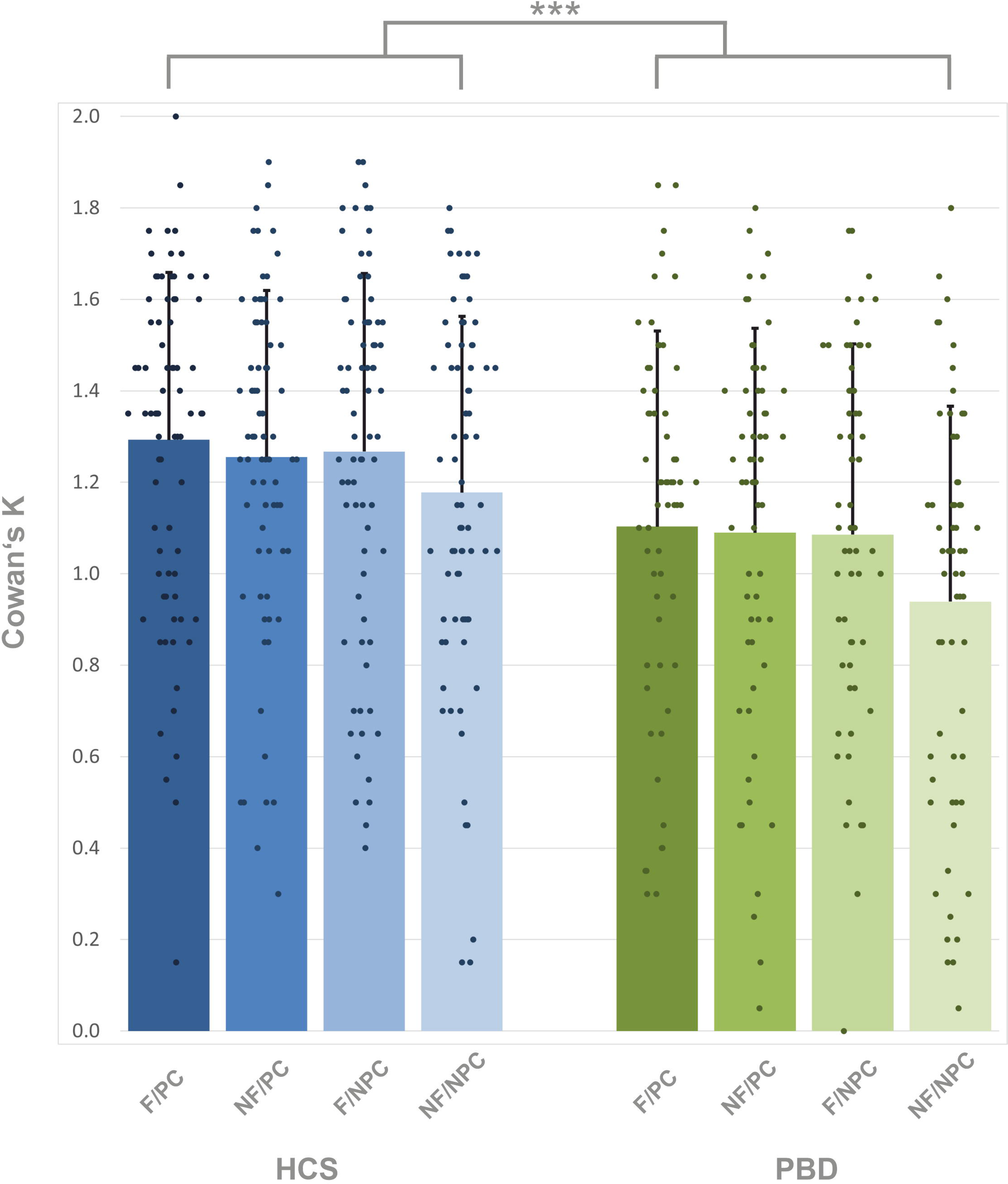
Amount of information stored in VWM in target trials, estimated with Cowan’s K in healthy control subjects = HCS and people with bipolar disorder = PBD. F/PC = flickering/predictive cue, NF/PC = non-flickering/predictive cue, F/NPC = flickering/non-predictive cue, NF/NPC = non-flickering/non-predictive cue. The between group contrast is displayed. *** indicates *p* < 0.001. Error bars indicate standard deviation.

**Table 2.** Overall significance estimates of the linear mixed model were obtained post-hoc with an ANOVA function using Satterthwaite’s method. Asterisks indicate significance *p* < 0.001 = ***, *p* < 0.01 = **, *p* < 0.05 = *.

There was no significant group difference for the efficiency of attentional prioritization (target Cowan’s K – catch Cowan’s K) in any condition. Yet, for the non-flickering/non-predictive cue condition there was a trend-level group difference (*p* = 0.057).

### EFFECTS OF SALIENCE AND CUE

For both groups we observed significant effects of salience as well as cue (both *p* < 0.001), but no significant two-way interactions of either group and salience, or group and cue. Furthermore, we did not observe a significant three-way interaction of group, salience and cue (Table 2). To confirm these findings, we conducted Bayes factor analyses for each effect, which provided moderate evidence in favor of an absence of effect (Table S4).

### ATTENTIONAL PRIORITIZATION AND VWM CAPACITY

The efficiency of attentional prioritization did not correlate with the independent measure of WM capacity (Pashler’s K) in any condition in PBD. It did however, correlate at a trend level, α = 0.05 in HCS in the non-flickering-bias/non-predictive cue condition (*p* = 0.046). However, the method by Steiger (Steiger, 1980) revealed no group difference for the strength of this correlation via a Fisher r to z transformation (z = 0.533, *p* = 0.297, Table 3).

**Table 3.** Results of Spearman correlations (two-tailed) of independent WM capacity estimate (Pashler’s K) and attentional prioritization efficiency (target Cowan’s K – catch Cowan’s K) within each condition, as well as mean Cowan’s K of each target condition. 95% confidence intervals are included, estimation of standard error is based on the formula proposed by Fieller, Hartley, and Pearson. Asterisks indicate significance *p* < 0.001 = ***, *p* < 0.01 = **, *p* < 0.05 = *. Abbreviations: HCS = healthy control subjects, PBD = people with bipolar disorder, CI = confidence level. The non-significant z-scores indicate no significant differences between groups of the correlations of VWM capacity and attentional prioritization in each condition.

### CORRELATION BETWEEN TARGET TRIALS AND VWM CAPACITY

VWM capacity correlated with Cowan’s K in each condition of target trials in HCS, yet in only three out of four conditions in PBD. There was no significant correlation between the flickering-bias/predictive cue condition and WM capacity. However, the Fisher r to z transformation revealed no group difference for the strength of this correlation (z = 0.453, *p* = 0.325, Table 3).

## DISCUSSION

We investigated the possible contribution of impaired attentional control to VWM dysfunction in BD. We manipulated the visual salience of targets and distractors as well as the predictiveness of external cues to isolate the impact of bottom-up and top-down attentional processing respectively. We observed a significant effect of group, with PBD storing less information in VWM across all conditions. Moreover, we observed a significant effect of salience that did not differ between groups, indicating a similar pattern of salience processing across groups. Furthermore, we observed a significant effect of cue that also did not differ between groups. Thus, the ability to utilize external top-down cues appears to be intact in both groups. However, it is important to note that no significant group interactions were observed, which would constitute evidence for group differences. The Bayes factor analysis revealed that these results were not attributable to a lack of statistical power. Consequently, these findings indicate that both top-down and bottom-up attention might be intact during VWM encoding in PBD (Mayer & Park, 2012).

Identifying specific cognitive processes that are impaired, and so-called ‘islands’ of preserved cognitive functioning provide important clues about the cognitive and neurophysiological mechanisms underlying attentional and VWM encoding dysfunction in neuropsychiatric disorders (Gold, Hahn, Strauss, & Waltz, 2009). Importantly, utilizing external cues did not improve performance of PBD up to the level of HCS. Thus, while our results do not implicate impaired attentional control, they do not fully refute the existence of a primary VWM encoding deficit in PBD. Based on the existing evidence for impairments of VWM encoding in PBD (Barnes-Scheufler et al., 2021; Huang et al., 2019; McKenna, Sutherland, Legenkaya, & Eyler, 2014), other aspects of this component process including VWM consolidation could account for the significant overall reduction in the amount of stored information in PBD. Moreover, disturbances during VWM maintenance and VWM retrieval could make additional contributions to the overall VWM dysfunction.

Our study confirms and extends the evidence for an overall WM deficit in PBD. These findings and the clear association between WM impairment and quality of life, functional outcome (Green, 2006) as well as psychosocial functioning (Iosifescu, 2012), underscores the relevance of this cognitive domain for cognitive remediation efforts including computerized training (Passarotti et al., 2020) in routine clinical practice (Miskowiak et al., 2018). Moreover, working memory is a key determinant of higher-order cognitive functions and intelligence (Hambrick, Kane, & Engle, 2005). Therefore, our results indicate that investigating the contribution of working memory dysfunction to general cognitive ability and to cognitive heterogeneity in ongoing large-scale studies of cognitive impairment in bipolar disorder (Burdick et al., 2019; Rabelo-da-Ponte et al., 2022) will be valuable to assess its overall impact on cognitive dysfunction in BD.

This is the first investigation to the best of our knowledge of the interaction of top-down and bottom-up attentional processes on VWM encoding in PBD. Contrary to our hypothesis, we observed no group difference in the efficiency of attentional prioritization. Analysis of catch trials revealed no significant effect of group (see supplementary materials). Thus, PBD did not store more goal-irrelevant information than HCS in any condition. The efficiency of attentional prioritization did not correlate with our independent measure of WM capacity in PBD and only as a trend in one condition in HCS. Conversely, our independent measure of WM capacity correlated with Cowan’s K in every target condition in HCS, but only in three out of four target conditions in PBD, yet Fisher r to z did not reveal any significant group differences.

Interestingly, the analysis of catch trials revealed a significant effect of cue (*p* = 0.001; Table S2), which we also observed in PSZ (Barnes-Scheufler et al., 2023). This suggests that PBD’s ability to use spatial cues to down-weight both salient and non-salient distractors is preserved. In light of the signal suppression hypothesis (Gaspelin & Luck, 2018b) this implies that when using external cues both PBD and HCS can sufficiently enhance top-down inhibitory control during attentional selection to increase local inhibition within early visual areas. However, our findings need to be interpreted with caution, considering the lack of group interactions. Alternatively, flickering might have had a negligible effect in the predictive-cue condition and cue properties might have had a negligible effect in the flickering condition. Moreover, given their chance-level performance in catch trials, both groups might have performed at ceiling level regarding top-down attention. This might have obscured subtle impairments of attention in PBD. Given the recognition of this domain as a key target of pro-cognitive treatment in PBD (Burdick et al., 2019; K. W. Miskowiak et al., 2017), further research into potential contribution of attentional dysfunction to VWM deficits is clearly warranted.

Abnormal GABAergic inhibitory neurotransmission has been proposed as a substrate of cognitive dysfunction across the schizo-bipolar spectrum (Prévot & Sibille, 2021; Volk & Lewis, 2014). Therefore, it is conceivable, that deficits in attention arise partly due to changes in GABAergic neurotransmission. Indeed, GABA levels appear to be reduced in PBD (Volk, Sampson, Zhang, Edelson, & Lewis, 2016), and in youth with BD cognitive deficits have been associated with GABA levels (Huber et al., 2018). However, extensive molecular phenotyping of GABAergic interneuron-related markers revealed that compared to PSZ, GABAergic abnormalities in PBD are less prevalent (Volk et al., 2016). In line with these studies, GABAergic deficits in PBD do not appear to impact the suppression of salient visual distractors to the same extent as in PSZ.

Functional neuroimaging has been used extensively to study the neurophysiological underpinnings of WM dysfunction in PBD. Common findings include a dysfunction of the dorsolateral prefrontal cortex (DLPFC) (Hamilton et al., 2009; Macoveanu et al., 2021; Miskowiak, Møller, & Ott, 2021; K. Miskowiak et al., 2017; Saldarini, Gottlieb, & Stokes, 2022; Thermenos et al., 2010) and the dorsal attention network comprising of the posterior parietal cortex (PPC) and frontal eye fields (FEF) (Brandt et al., 2014). Furthermore, there have also been reports of decreased activation in the left lateral occipital cortex, part of the ventral visual stream, implicating impaired visual processing in VWM dysfunction (Manelis et al., 2022). However, studies employing paradigms designed to distinguish between the major VWM component processes remain more limited. This approach revealed BD hypoactivation in the PPC during the encoding interval, with subsequent hypoactivation in visual areas during the maintenance interval (Huang et al., 2019). This finding is particularly noteworthy because of the close relationship between activation in these brain areas during encoding and maintenance with VWM capacity (Linden et al., 2003; Todd & Marois, 2004). While these areas are also closely linked to top-down attention during VWM encoding (Mayer et al., 2007), based on our current results we would expect to observe a specific link of abnormal activation in these areas with impaired VWM performance.

Importantly, our findings differ in crucial ways from our previous results in PSZ (Barnes-Scheufler et al., 2023). While we did not conduct any direct statistical comparisons between both data sets, several inferences can still be made. Both PSZ and PBD stored significantly less information across all conditions compared to HCS. However, in contrast to our study in PSZ, we did not observe group by salience and/or group by cue effects in PBD. Thus, the pattern of cognitive impairment revealed by our paradigm in PBD appears to differ from PSZ.

We consider the relatively large sample size of patients encompassing bipolar types 1 and 2 both with and without a history of psychosis but without any psychiatric comorbidities a strength of our study. However, a notable limitation pertains to the design of our paradigm, which might have prevented the detection of subtle attentional deficits For instance, there is evidence indicating that cue-driven attention allocation might not necessarily be fully endogenous (Tipples, 2002), which might limit the ability to fully separate bottom-up and top-down attentional processes. Due to the use of stimulus orientation rather than location as memoranda, automatic attentional capture of the locations of distractors might not have sufficiently facilitated the encoding of their orientation. Encoding this additional stimulus feature despite its low relevance would have likely required the voluntary allocation of additional attentional resources. This might have obscured possible deficits in the down-weighting of salient distractors in PBD. Additionally, our sample of PBD had a relatively high mean IQ, which is not fully representative of the entire population and could have been associated with better performance. Lastly, we did not record the overall number of mood episodes in PBD, which would have allowed us to explore the impact of illness severity on visual WM dysfunction (Rabelo-da-Ponte et al., 2022).

To summarize, we observed a significant reduction in the amount of information encoded in PBD across all conditions. We also provide evidence for preserved cognitive processes in BD namely the successful utilization external top-down cues and stimulus salience during WM encoding. Our results underscore the importance of behavioral paradigms geared toward measuring cognitive constructs derived from cognitive neuroscience to identify impaired and intact component processes of WM and attention in PBD.

## ACKNOWLEDGEMENTS

The authors are very thankful to Prof. Dr. Christoph Fehr, Dr. Peter Hustedt and Hannah Schroeder for their support in recruiting patients, and Tobias Lehmann and Caroline Mirkes for assisting with data acquisition.

## AUTHORS’ CONTRIBUTIONS

All authors made substantial contributions to the conception or design of the work, or the acquisition, analysis or interpretation of data. Authors LR, MS, BP, JSM, MW, and RAB designed the experiment. Authors CVB-S and LR acquired the data. Authors CVB-S, CS, JSM, SM and RAB analyzed the data. Authors CVB-S, SK-S and RAB undertook the literature searches and wrote the first draft of the manuscript. All authors contributed to and revised the manuscript. All authors read and approved the final manuscript.

## PUBLICATION HISTORY

This manuscript was previously posted to bioRxiv: https://doi.org/10.1101/2024.01.04.574011

## FUNDING STATEMENT

We report no financial relationships with commercial interests. C.V Barnes-Scheufler was supported by a “main doctus” scholarship from The Polytechnic Foundation of Frankfurt am Main.

## CONFLICTS OF INTEREST

The Authors have declared that there are no conflicts of interest in relation to the subject of this study.

## ETHICAL STANDARDS

The authors assert that all procedures contributing to this work comply with the ethical standards of the relevant national and institutional committees on human experimentation and with the Helsinki Declaration of 1975, as revised in 2008. The ethics committee of the University Hospital Frankfurt approved all study procedures (experiment number 2/15).

## Notes

### Competing Interest Statement

The authors have declared no competing interest.

### Summary of Updates

Major revision of all aspects of the manuscript.

