## Supplementary material for "Impaired Visuospatial Working Memory but Preserved Attentional Control in Bipolar Disorder": Tables

Table 1. Participant characteristics of both groups including information regarding demographics, psychopathology and current medication.

| | HCS (n = 76) | | PBD (n = 63) | | Statistic | df | *p* |
|---|---|---|---|---|---|---|---|
| **Demographics** | **Mean** | **SD** | **Mean** | **SD** | | | |
| Age, Years | 39.01; range 19-61 | 12.96 | 42.29; range 20-61 | 11.06 | $t$ = 1.58 | 137 | 0.116 |
| Handedness | 69.82 | 37.30 | 64.83 | 11.06 | $t$ = -0.69 | 137 | 0.490 |
| Participant Education, Years | 16.09 | 2.16 | 16.51 | 2.23 | $t$ = 1.15 | 137 | 0.267 |
| Parental Education, Years | 14.61 | 3.06 | 14.90 | 3.60 | $t$ = 0.53 | 136 | 0.598 |
| Premorbid IQ | 115.29 | 12.51 | 117.60 | 12.21 | $t$ = 1.09 | 136 | 0.278 |
| Duration of Illness, Years | | | 12.32; range 1-35 | 8.89 | | | |
| Age of Illness Onset, Years | | | 29.97; range 9-50 | 11.83 | | | |
| Sex, Male/Female | 27 / 49 | | 16 / 47 | | $X^2$ = 1.65 | 1 | 0.198 |
| PBD I | | | 38 | | | | |
| PBD II | | | 25 | | | | |
| PBD HPS+ | | | 32 | | | | |
| PBD HPS- | | | 31 | | | | |
| **Psychopathology** | | | **Mean** | **SD** | | | |
| YMRS | | | 1.40 | 1.73 | | | |
| MADRS | | | 3.03 | 2.70 | | | |
| **Medication** | | | | | | | |
| SGA | 0 | | 31 (49%) | | | | |
| Lithium | 0 | | 29 (46%) | | | | |
| Valproate | 0 | | 11 (17%) | | | | |
| Lamotrigine | 0 | | 12 (19%) | | | | |
| Antidepressants | 0 | | 22 (35%) | | | | |

Table 2. Results of the linear mixed model investigating the effects of cue (predictive cue/non-predictive cue) and salience (flickering/non-flickering) in target trials in both PBD and HCS with the covariate of age.

| | **df** | **Sum of Squares** | **Mean Square** | **F** | ***p*** |
|---|---|---|---|---|---|
| Group | 139 | 0.425 | 0.425 | 8.140 | 0.005** |
| Salience | 417 | 0.711 | 0.711 | 13.623 | < 0.001*** |
| Cue | 417 | 0.635 | 0.635 | 12.170 | < 0.001*** |
| Age | 139 | 1.124 | 1.124 | 21.546 | < 0.001*** |
| Group x Salience | 417 | 0.010 | 0.010 | 0.184 | 0.669 |
| Group x Cue | 417 | 0.036 | 0.036 | 0.697 | 0.404 |
| Salience x Cue | 417 | 0.296 | 0.296 | 5.668 | 0.018* |
| Group x Salience x Cue | 417 | 0.057 | 0.057 | 1.093 | 0.297 |

Table 3. Correlational analyses with working memory capacity

| **Attentional prioritization and visual working memory capacity** | **Statistic** | ***p*** | ***CI lower*** | ***CI upper*** |
|---|---|---|---|---|
| **HCS (*n* = 76 )** | | | | |
| Flickering-bias/predictive cue | $r_s$ = 0.137 | 0.238 | -0.098 | 0.358 |
| Flickering-bias/non-predictive cue | $r_s$ = 0.203 | 0.079 | -0.030 | 0.415 |
| Non-flickering-bias/predictive cue | $r_s$ = 0.149 | 0.199 | -0.086 | 0.386 |
| Non-flickering-bias/non-predictive cue | $r_s$ = 0.230 | 0.046* | -0.002 | 0.438 |
| **PBD (*n* = 62)** | | | | |
| Flickering-bias/predictive cue | $r_s$ = 0.139 | 0.281 | -0.122 | 0.382 |
| Flickering-bias/non-predictive cue | $r_s$ = 0.088 | 0.497 | -0.173 | 0.337 |
| Non-flickering-bias/predictive cue | $r_s$ = 0.148 | 0.252 | -0.113 | 0.390 |
| Non-flickering-bias/non-predictive cue | $r_s$ = 0.140 | 0.276 | -0.121 | 0.383 |
| **Fisher z-transformation** | | | | |

| | | | | |
|---|---|---|---|---|
| Flickering-bias/predictive cue | *z* = 0.012 | 0.495 | | |
| Flickering-bias/non-predictive cue | *z* = 0.672 | 0.251 | | |
| Non-flickering-bias/predictive cue | *z* = 0.006 | 0.498 | | |
| Non-flickering-bias/non-predictive cue | *z* = 0.533 | 0.297 | | |
| **Target trials and visual working memory capacity** | | | | |
| **HCS (*n* = 76 )** | | | | |
| Flickering-bias/predictive cue | $r_s$ = 0.271 | 0.018* | 0.042 | 0.473 |
| Flickering-bias/non-predictive cue | $r_s$ = 0.500 | < 0.001*** | 0.303 | 0.656 |
| Non-flickering-bias/predictive cue | $r_s$ = 0.370 | 0.001** | 0.151 | 0.554 |
| Non-flickering-bias/non-predictive cue | $r_s$ = 0.317 | 0.005** | 0.092 | 0.511 |
| **PBD (*n* = 62)** | | | | |
| Flickering-bias/predictive cue | $r_s$ = 0.196 | 0.126 | -0.064 | 0.431 |
| Flickering-bias/non-predictive cue | $r_s$ = 0.329 | 0.009** | 0.079 | 0.540 |
| Non-flickering-bias/predictive cue | $r_s$ = 0.268 | 0.036* | 0.012 | 0.491 |
| Non-flickering-bias/non-predictive cue | $r_s$ = 0.312 | 0.013* | 0.060 | 0.527 |
| **Fisher z-transformation** | | | | |
| Flickering-bias/predictive cue | *z* = 0.453 | 0.325 | | |
| Flickering-bias/non-predictive cue | *z* = 1.186 | 0.118 | | |
| Non-flickering-bias/predictive cue | *z* = 0.650 | 0.258 | | |
| Non-flickering-bias/non-predictive cue | *z* = 0.032 | 0.487 | | |
