## Supplemental_material for "Impaired Visuospatial Working Memory but Preserved Attentional Control in Bipolar Disorder"

### METHODS:

*Main working memory task:*

HCS were screened using the Structural Clinical Interview SCID-I from the Diagnostic and Statistical Manual, Version IV, in German (Saß, Wittchen, Zaudig, & Houben, 2003).

Participants were informed about the currently task-relevant Gabor patches (flickering-bias or non-flickering-bias) and of the high likelihood that they would be probed during retrieval. For example, in the flickering-bias predictive cue condition, the instructions read ‘In this task, the flickering-bias patterns will be probed preferentially. The positions of these striped patterns will be marked by means of the fixation cross,’ and was displayed for 15 seconds.

Because a partial display was used at recall, we calculated the amount of information stored in VWM using Cowan’s K instead of Pashler’s K (Rouder, Morey, Morey, & Cowan, 2011).

Post-hoc contrasts of the LMM were obtained with the emmeans package (Russell V Lenth, 2023).

*Investigation of overall possible influences of medication*

We were interested in investigating possible influences of medication on the amount of information store into WM. To this end, we performed Spearman bivariate correlations (two-tailed) in PBD of daily dose in mg of lithium, valproate and lamotrigine with overall target Cowan’s K. In addition, we conducted a t-test (2-tailed) to check for a possible difference between PBD receiving antipsychotic medication at the time of the study and those PBD that were not receiving antipsychotic medication.

*Group differences in attentional prioritization*

We compared attentional prioritization (target Cowan’s K – catch Cowan’s K) in each condition across groups using independent sample t-tests (Table S1).

*Catch trials*

In 20% of trials a non-target Gabor patch was probed at retrieval- for example when the instructions indicated that flickering information should be encoded (‘flickering-bias’, a non-flickering Gabor patch was probed (Figure 1). If a predictive cue was displayed, that cue indicated the incorrect locations. Overall accuracy across all four catch conditions was around chance level (PBD = 56%, HCS = 56%).

We conducted an LMM using catch trials to predict the WM score with group, salience, cue and age (Formula: score ~ Salience * Cue * Group + Age). The model included subject as a random effect (formula: ~1 | Subject), (Table S2).

The post-hoc contrasts were computed using the Kenward-Roger degrees-of-freedom method with pairwise t-tests adjusted with the Tukey method and were adjusted for age. We investigated differences between and within groups (Table S3).

*Bayes Factor Analyses*

Bayes factor analyses of all main effects were conducted (Table S4).

*Investigation of the impact of duration of illness*

Duration of illness was operationalized as years since the diagnosis of bipolar disorder. We conducted a repeated-measures ANOVA within the PBD group with the factors of salience and cue and the covariates of age and duration of illness (Table S5).

*Working memory capacity task*

We implemented a ‘canonical’ color change detection task (Figure S1) on a personal computer using Presentation software in Version 14.9 (www.neuro bs.com). Stimuli were presented on a grey background (RGB values: 191, 191, 191) in a dimly lit room with a viewing distance of approximately 60 cm. Throughout the experiment, a black fixation cross was displayed at the center of the screen. Each trial began with the alert phase, during which the fixation cross turned to red for 500 ms. This was followed by a preparation phase of 500 ms. During the encoding phase a sample array of four colored circles was presented for 200 ms. Each circle had a visual angle of approximately 0.95°. These circles were spaced equally apart on an imaginary circle with 12 possible locations around the black fixation cross covering a visual angle of approximately 5.25°, and the minimum distance between two circles was 0.29°. Each circle had one of seven easily discriminable possible colors with the following RGB values: black (0, 0, 0), red (255, 0, 0), white (255, 255, 255), blue (0,

0, 255), green (0, 255, 0), yellow (255, 255, 0), and magenta (255, 0, 255), with no repetitions of colors within a trial. During the delay phase, the black fixation cross remained on the screen for 1800 ms. A whole-display recognition test array followed, in which participants had a maximum duration of 3000 ms to decide if the test array was identical to the sample array presented in the encoding phase, or if one of the circles had changed color. Because a whole-display recognition was used, we calculated the amount of information stored in WM using Pashler's K. Half of the trials were change trials (right mouse button), the other half no-change trials (left mouse button). In change trials, a randomly chosen circle changed its color. The total duration of each trial was 6000 ms followed by an inter-trial interval of 3000 ms. All participants received the same instructions prior to the beginning the task, and were asked to perform as accurately as possible, and to keep their eyes fixated constantly on the center of the screen. A total of 60 trials were tested in each participant, which required approximately nine minutes of testing time (Barnes-Scheufler et al., 2021).

### Figure S1. Working Memory Capacity Task

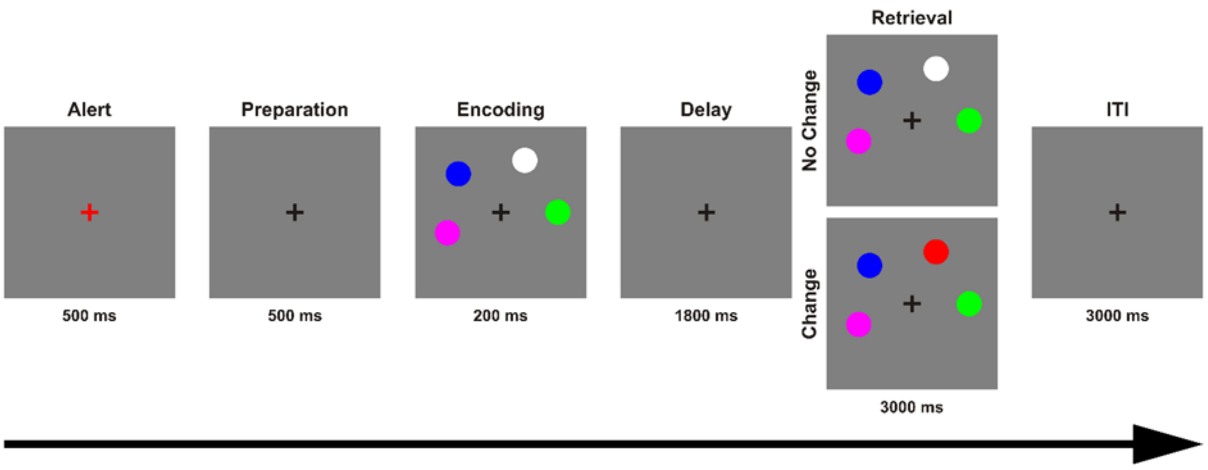

Figure S1. The change detection task used to assess working memory capacity. Each trial began with the alert phase, during which the fixation cross turned to red for 500 ms. This was followed by a preparation phase of 500 ms. During the encoding phase a sample array of four colored circles was presented for 200 ms. During the delay phase, the black fixation cross remained on the screen for 1800 ms. The whole-display recognition test array followed, in which participants had a maximum duration of 3000 ms to decide if the test array was identical to the sample array presented in the encoding phase, or if one of the circles had changed color. See (Barnes-Scheufler et al., 2021) for more details.

### RESULTS:

*Investigation of overall possible influences of medication*

There were no significant correlations between daily dose of lithium ($r_s$ = 0.178, $p$ = 0.355), valproate ($r_s$ = -0.239, $p$ = 0.479), or lamotrigine ($r_s$ = -0.413, $p$ = 0.207) with overall target Cowan's K in PBD. There was no significant difference in overall target Cowan's K between PBD who were taking antipsychotics (n = 31), and those who were not (n = 32, t(61) = 0.593, $p$ = 0.556).

*Investigation of the impact of duration of illness*

There was no significant effect of duration of illness ($p$ = 0.742) within PBD, and therefore we did not include this covariate in our main LMM analysis.

### Table S1. Group differences in attentional prioritization

| | *t* | *df* | *p Value* |
|---|---|---|---|
| Flickering-bias/predictive cue | -1.529 | 137 | 0.128 |
| Flickering-bias/non-predictive cue | -1.421 | 137 | 0.158 |
| Non-flickering-bias/predictive cue | -1.779 | 137 | 0.077 |
| Non-flickering-bias/non-predictive cue | -1.920 | 137 | 0.057 |

Table S1: Results of independent sample t-tests. We did not observe any significant difference between groups in attentional prioritization.

### Figure S2. Results of catch trials

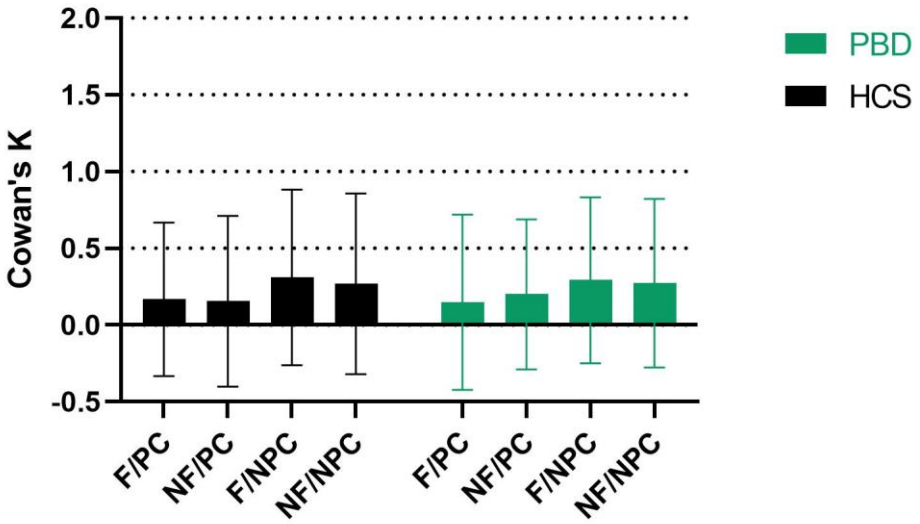

Figure S2. Amount of information stored in VWM in catch trials, estimated with Cowan's K in healthy control subjects = HCS and people with bipolar disorder = PBD. F/PC = flickering-bias/predictive cue, NF/PC = non-flickering-bias/predictive cue, F/NPC = flickering-bias/non-predictive cue, NF/NPC = non-flickering-bias/non-predictive cue. Error bars indicate standard deviation. We observed solely a significant effect of cue in the catch trials ($p$ = 0.001). There were no significant effects of group, salience, or age (Table S2). Furthermore, there were no significant interactions of group by salience, group by cue, salience by cue, and no three-way interaction of group by salience by cue.

### Table S2. Results of LMM for catch trials

| | *df* | *F* | *p Value* |
|---|---|---|---|
| Group | 139 | 0.02 | 0.880 |
| Salience | 417 | 0.03 | 0.873 |
| Cue | 417 | 10.32 | 0.001* |
| Age | 139 | 1.03 | 0.312 |
| Group x Salience | 417 | 0.35 | 0.553 |
| Group x Cue | 417 | 0.07 | 0.788 |
| Salience x Cue | 417 | 0.45 | 0.501 |
| Group x Salience x Cue | 417 | 0.08 | 0.781 |

Table S2: Results of LMM of catch trials with factors salience and cue, and covariate age. We observed solely a significant effect of cue in the catch trials. Asterisks indicate significance $p < 0.001$ = ***, $p < 0.01$ = **, $p < 0.05$ = *.

### Table S3. Results of between and within group comparisons of catch trials

| | *t* | *df* | *p Value* |
|---|---|---|---|
| **Between Group (*n* = 139)** | | | |
| Flickering-bias/predictive cue | 0.125 | 400 | 0.901 |
| Flickering-bias/non-predictive cue | 0.117 | 400 | 0.907 |
| Non-flickering-bias/predictive cue | -0.555 | 400 | 0.579 |
| Non-flickering-bias/non-predictive cue | -0.128 | 400 | 0.898 |
| **HCS (*n* = 76 )** | | | |
| Flickering-bias/predictive cue vs Flickering/non-predictive cue | -2.02 | 423 | 0.182 |
| Flickering-bias/predictive cue vs Non-flickering/predictive cue | 0.187 | 423 | 0.998 |
| Non-flickering-bias/predictive cue vs Non-flickering/non-predictive cue | -1.609 | 423 | 0.375 |
| Non-flickering-bias/non-predictive cue vs. Flickering/non-predictive cue | -0.599 | 423 | 0.933 |
| **PBD (*n* = 63)** | | | |
| Flickering-bias/predictive cue vs Flickering/non-predictive cue | -1.849 | 423 | 0.252 |
| Flickering-bias/predictive cue vs Non-flickering/predictive cue | -0.657 | 423 | 0.913 |
| Non-flickering-bias/predictive cue vs Non-flickering/non-predictive cue | -0.945 | 423 | 0.781 |
| Non-flickering-bias/non-predictive cue vs. Flickering/non-predictive cue | -0.247 | 423 | 0.995 |

Table S3: Results of post-hoc t-tests between and within groups between conditions in catch trials. We did not observe any significant differences between groups in the catch trial conditions or between conditions within groups.

### Table S4. Results of Bayes factor analyses of main effects

| Term | Against Model | BF10 (Bayes Factor) | CI ± (%) | Interpretation |
|---|---|---|---|---|
| **Full Model** | Intercept only | 6.755764e+13 | ±3.86% | Very strong evidence for full model |
| **Group** | Score ~ Cue + Salience + Group * Cue + Group * Salience + Salience * Cue + Group * Salience * Cue + Age | 1 | ±5.46% | No evidence |
| **Salience** | Score ~ Cue + Group + Group * Cue + Group * Salience + Salience * Cue + Group * Salience * Cue + Age | 0.995977 | ±5.54% | No evidence |
| **Cue** | Score ~ Salience + Group + Group * Cue + Group * Salience + Salience * Cue + Group * Salience * Cue + Age | 0.996056 | ±5.53% | No evidence |
| **Age** | Score ~ Cue + Salience + Group + Group * Cue + Group * Salience + Salience * Cue + Group * Salience * Cue | 3.61323e+11 | ±5.63% | Very strong evidence |
| **Cue:Salience** | Score ~ Cue + Salience + Group + Group * Cue + Group * Salience + Group * Salience * Cue + Age | 1 | ±5.46% | No evidence |
| **Cue:Group** | Score ~ Salience + Group + Group * Salience + Salience * Cue + Group * Salience * Cue + Age | 1.000238 | ±5.45% | No evidence |
| **Salience:Group** | Score ~ Cue + Salience + Group + Group * Cue + Salience * Cue + Group * Salience * Cue + Age | 1.000264 | ±5.45% | No evidence |

| | | | | |
|---|---|---|---|---|
| **Group:Salience:Cue** | Score ~ Cue + Salience + Group + Group * Cue + Group * Salience + Salience * Cue + Age | 0.1865021 | ±5.42% | Strong evidence for absence of interaction |

#### Table S5. Results of ANOVA in PBD for covariate duration of illness

| | **df** | **Sum of Squares** | **Mean Square** | **F** | ***p*** |
|---|---|---|---|---|---|
| Age | 1 | 2.92 | 2.92 | 5.184 | 0.026* |
| Duration of illness | 1 | 0.06 | 0.06 | 0.109 | 0.742 |
